# Dashboard-style interactive plots for RNA-seq analysis are R Markdown ready with *Glimma* 2.0

**DOI:** 10.1101/2021.07.30.454464

**Authors:** Hasaru Kariyawasam, Shian Su, Oliver Voogd, Matthew E. Ritchie, Charity W. Law

## Abstract

*Glimma* 1.0 introduced intuitive, point-and-click interactive graphics for differential gene expression analysis. Here, we present a major update to *Glimma* which brings improved interactivity and reproducibility using high-level visualisation frameworks for R and JavaScript. *Glimma* 2.0 plots are now readily embeddable in R Markdown, thus allowing users to create reproducible reports containing interactive graphics. The revamped multidimensional scaling plot features dashboard-style controls allowing the user to dynamically change the colour, shape and size of sample points according to different experimental conditions. Interactivity was enhanced in the MA-style plot for comparing differences to average expression, which now supports selecting multiple genes, export options to PNG, SVG or CSV formats and includes a new volcano plot function. Feature-rich and user-friendly, *Glimma* makes exploring data for gene expression analysis more accessible and intuitive and is available on Bioconductor and GitHub.

## Introduction

RNA-sequencing (RNA-seq) is a high-throughput method for characterising transcriptomes (1). Researchers commonly leverage RNA-seq technology to compare the transcription levels of genes across experimental conditions, in a workflow known as differential expression (DE) analysis. As there are tens of thousands of genes involved in DE analyses, it can be difficult to pinpoint information on genes of interest in densely populated static R (2) plots. Static graphics necessarily provide a flattened perspective on the data; version 1 of *Glimma* (3) aimed to remedy this by allowing users to interactively explore data at the sample-level through dimensionality reduction plots and at the gene-level in plots of summary statistics obtained from popular DE analysis tools *limma* (4), *edgeR* (5) and *DESeq2* (6).

While many powerful tools exist for producing interactive plots for DE analysis (7–11), they require users to run a *Shiny* server (12), which may be difficult to navigate for those with minimal experience in R. The *Glimma* software does not depend on *Shiny* and can produce portable outputs that can be viewed without an active installation of R. This allows the package to cater to its main user base of biologists and end-users who would like drop-in graphics for popular gene expression workflows.

*Glimma* version 1 allowed the creation of two interactive versions of *limma*-style plots: a multidimensional scaling (MDS) plot used to assess variability between samples, and a mean-difference (MD) plot used for identifying differentially expressed genes between experimental conditions. These could be exported as HTML files and shared with collaborators, allowing biologists to investigate interesting features in the data with minimal coding required.

The responsive and user-friendly layout of *Glimma* 1.0 has proven very popular among the Bioconductor community, amassing over 19,000 downloads in 2020 alone. The software is commonly used for exploration of transcriptional data from raw gene expression-levels to summarised results obtained from DE analysis (13).

Intended as a drop-in interactive visualisation tool for common RNA-seq workflows, *Glimma* was built on d3.js (14) and relied on custom-built functions for connecting R code with a web-based frontend. The low-level nature of d3.js and the high complexity of the codebase made it difficult to add improved interactivity and new output formats to the package. For instance, simple improvements such as adding plot legends and scaling point sizes became intractable tasks. Version 2.0 of *Glimma* was built to address these deficiencies by reproducing all existing functionality in 1.0 using existing high-level libraries such as *htmlwidgets* (15) and *Vega* (16). This would make it easier for bioinformaticians and software engineers to rapidly develop new features in response to new developments in gene expression studies, such as single cell RNA-seq (scRNA-seq) analysis.

Another important theme in the second iteration of *Glimma* was reproducibility. Previous versions instantiated plots within a new HTML window when *Glimma* R functions were called, creating a separation between interactive plots and the code that created them. It was therefore a requirement that plots could be embedded in R markdown (17) in version 2, allowing bioinformaticians to share reports with graphics interwoven between blocks of code.

*Glimma* 2.0 offers three main interactive plots - an MDS plot via the `glimmaMDS` function, an MD plot (which has been renamed to MA plot) via the `glimmaMA` function, and a volcano plot via the `glimmaVolcano` function. Additionally, a generalised version of an interactive plot that comprises of the two components seen in the MD and volcano plots, which includes a plot of summary results from a DE analysis and a plot of expression data from which the DE analysis was performed on, is available via the `glimmaXY` function (see Figure 1). We refer to the style of these plots as “summary-expression” plots for the remainder of the article.

**Fig. 1.**
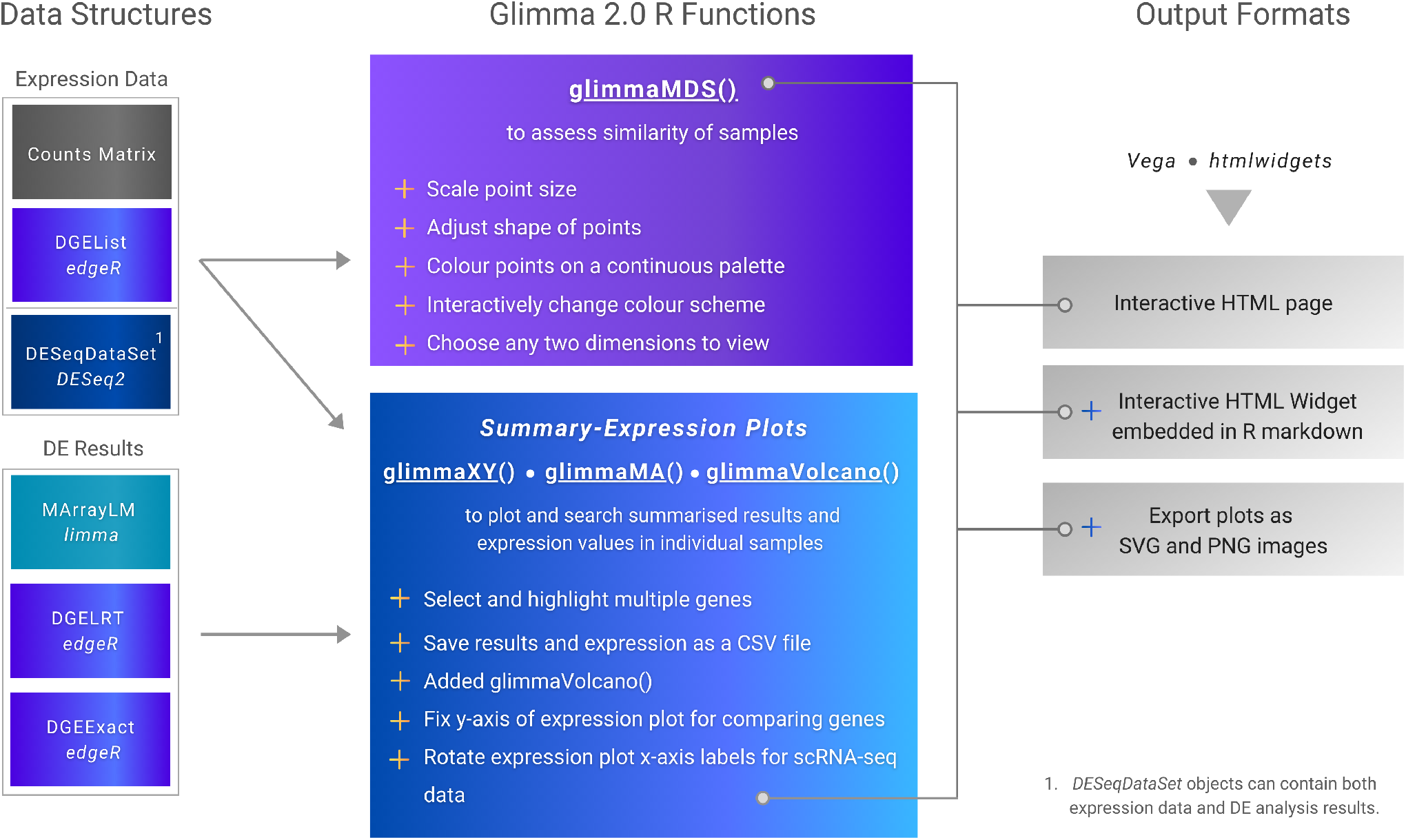
Schematic diagram for *Glimma* 2.0. Input data structures from *limma, edgeR* and *DESeq2* RNA-seq analysis workflows are shown on the left, connected to the *Glimma* functions that accept them. New features for the MDS and summary-expression plots (MA, volcano and XY) are shown in the centre, demarcated by plus signs. Function outputs are listed on the right with several new output formats shown.

Relative to the original version of the software, the volcano plot is a new addition to *Glimma*. It was included for ease of use since this it is a common plot created in a RNA-seq workflow. The new plot simplifies coding by *Glimma*-users who would have otherwise had to used the more general XY plot to create this plot.

The MD plot has been renamed to “MA plot” to reflect the original name given to this style of plot in the *limma* package. A plot of mean gene expression (or averages) against log_2_-fold change or logFC (also called “M-values”) were referred to as an MA plot. *Glimma*’s MA plot is equivalent to *limma*’s MD plot.

In this article, we will demonstrate a suite of interactivity and functionality upgrades made in *Glimma* 2.0. The embedded plots offer a seamless user interface integrated with the workflow of DE analysis. Amidst fervent demand for improved reproducibility in bioinformatics research, we anticipate that version 2 will be even more popular than the initial version.

## MATERIALS AND METHODS

### Availability

Functions from *Glimma* 1.0, as well as the improvements made to *Glimma* 2.0 are available in the *Glimma* software package. The stable version of the software is released on Bioconductor (18), and is available at https://bioconductor.org/packages/Glimma. Both stable and developmental releases of *Glimma* are available on GitHub at https://github.com/hasaru-k/GlimmaV2.

### Frameworks

*Glimma* 2.0 was re-developed using *htmlwidgets* for R (15), a framework for creating web-based visualisations which behave like R plots, which offers embeddability in R Markdown out of the box, addressing reproducibility issues in the previous version. Some popular packages using *htmlwidgets* as a foundation include *Plotly* (19) and *dygraphs* for R (20). The *htmlwidgets* package provides a convenient API (application programming interface) for software developers to bind R functions to a JavaScript visualisation front-end codebase. This means the process of connecting R function calls to web-based visualisations including serialisation of R data structures is automated, greatly reducing the complexity of the development process. To ensure backwards compatibility with previous versions, version 2 was built to support data structures from *edgeR, limma* and *DESeq2* workflows as inputs (see Figure 1).

### Visualisation

Interactive plots were generated within an instantiated widget using the *Vega* visualisation grammar (16) which is based on d3.js (14) but provides higher level graphical primitives and allows convenient linkage of graphs. Declarative JSON specifications for each genre of plot were created, parameterised and passed to the *Vega* API which renders the desired interactive web-based graphics. The *Vega* API was also used to display tooltips, respond to on-click events and dynamically update plots.

In conjunction with external libraries, native JavaScript was used to link data across various plots and tables. In the front-end interface for summary statistic plots, an interaction state machine ensures that the response to any sequence of inputs from the user is well-defined. For instance, selecting genes on the summary plot triggers a transition into the graph selection state. This event filters the table to display the selected genes and updates other visual controls. The open-source *FileSaver*.*js* package (21) was used for exporting gene summary statistics and expression data as CSV files for summary-expression plots.

### Demonstration Dataset

A dataset from Sheridan *et al*. (2015) (22) was utilised to generate the *Glimma* 2.0 plots illustrated in Figures 2 and 3. The experiment includes three cell populations: basal, luminal progenitor (LP) and mature luminal (ML) cells from the mammary glands of female mice. Three RNA samples were sequenced from each cell population, resulting in nine samples in total. Read alignment to the *mm10* mouse reference genome was performed using the *Rsubread* package (23) (version 1.14.1) to obtain gene-level counts. Expression data and details regarding experimental design are available from [Gene Expression Omnibus](https://www.ncbi.nlm.nih.gov/geo/) under accession number GSE63310. Interactive versions of Figures 2 and 3 can also be accessed through the *Glimma* 2.0 vignettes.

**Fig. 2.**
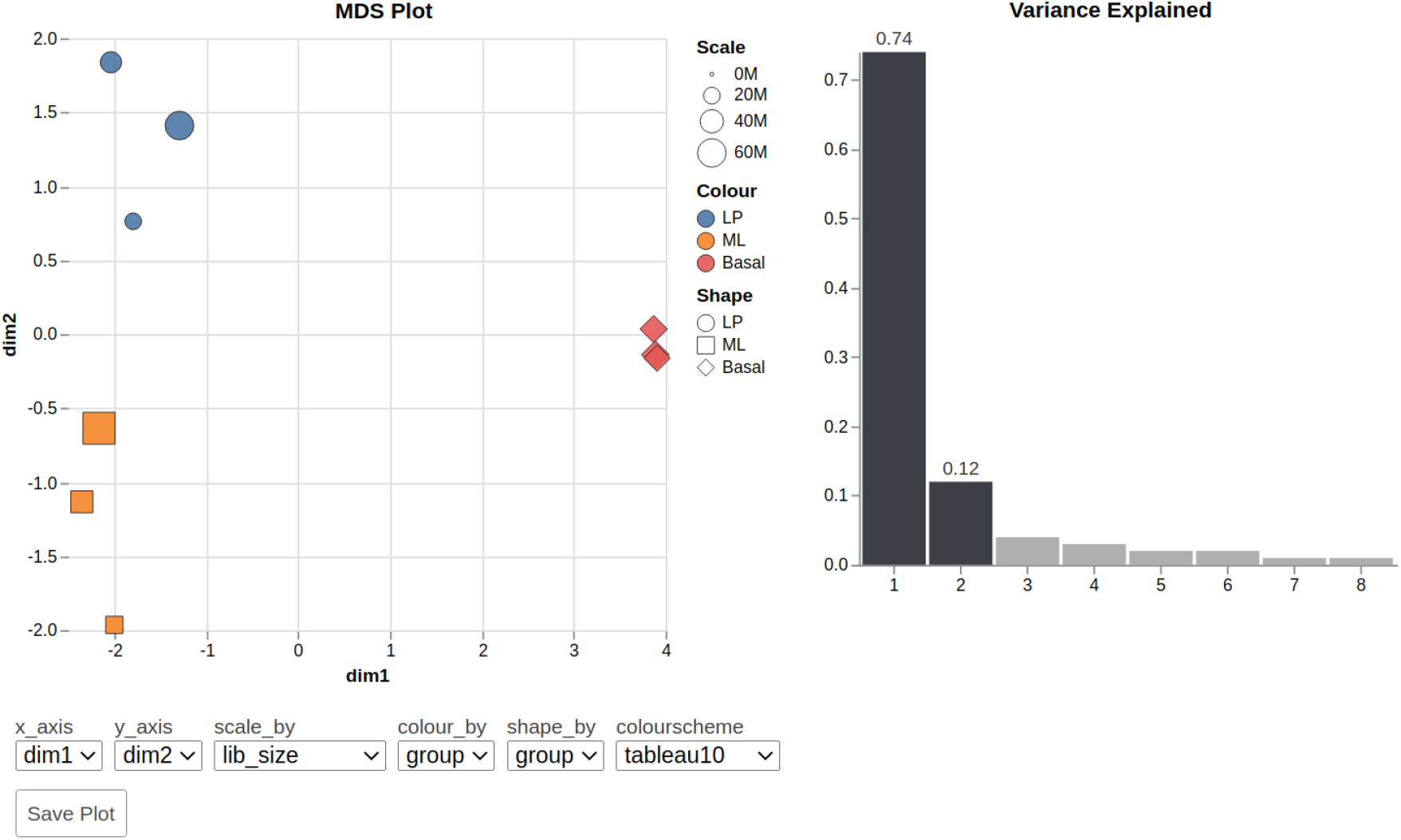
Screenshot of a markdown-embedded *Glimma* 2.0 MDS plot with the first and second dimensions on display. Each sample point is coloured and shaped according to the experimental group and scaled by the library size. The bar plot on the right displays the proportion of variance accounted for by each dimension. The drop-down boxes allow users to view the combination of any two dimensions, and change the attribute for which points are scaled, coloured and shaped using a selection of colour schemes. The plots can be saved as a static plot for presentations and publication using the “Save Plot” button.

**Fig. 3.**
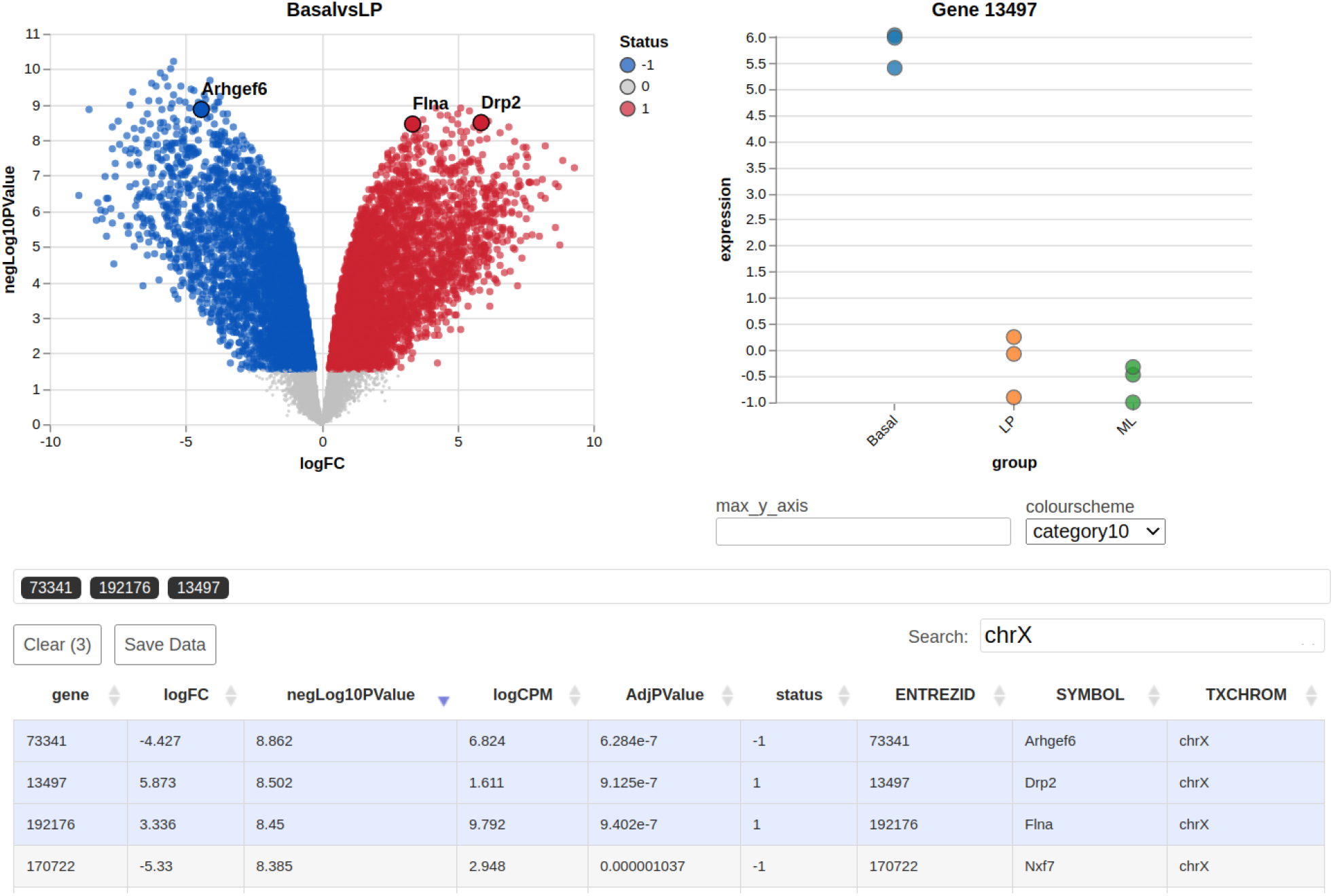
Screenshot of a markdown-embedded *Glimma* 2.0 volcano plot displaying gene-wise logFC on the x-axis against *−* log_10_(*p − value*) on the y-axis. The three most significant genes on the X chromosome are highlighted. Hovering over points in the summary plot displays tabular details about the gene. The expression plot for the most recently selected gene, *Drp2* with Entrez gene identifier 13497, is shown on the right with log-CPM plotted against experimental groups. Hovering over points in the expression plot displays the sample name and expression value. The filtered table beneath provides full summary statistics and annotations for the selected genes.

## RESULTS

### Embedded Plot or Stand-Alone File

A major new feature of *Glimma* 2.0 is that it allows plots to be embedded in R markdown. Users still have the option to export widgets as a standalone HTML file with inlined JavaScript, CSS and assets. This is another improvement on version 1.0 where exported HTML had to be manually zipped with scripts and assets before sharing interactive plots with collaborators.

### Image Export

*Glimma* 2.0 widgets contain save buttons which allow the user to export plots in their current state as static PNG or SVG files. The latter filetype is an infinitely scalable vector format which allows users to further manipulate plots in Adobe Illustrator. For MDS plots, both the MDS plot itself and the “Variance Explained” plot can be exported separately using the “Save Plot” button. Similarly, summary-expression plots have the capability to export either of the summary or expression plots separately. This is a marked improvement over version 1.0 which lacked image saving capabilities.

### MDS Plot Improvements

Each point on the MDS plot represents an experimental sample for which the transcriptional output across thousands of genes has been recorded. MDS reduces the dimensionality of the data so that the first few dimensions represent most of the variation, which allows us to visually examine similarities and differences in the transcription profiles of samples (4). Samples that cluster together are interpreted as more similar than those which are far apart, and each dimension of the MDS plot explains a decreasing proportion of the total variation. This is used for exploratory analysis of an experiment to determine which factors drive the variation between samples and the degree of their effects. It is useful for observing both expected and unexpected differences in the samples’ transcriptional profiles (e.g. separation between biological groups versus unknown batch effects), and serves as a predictor of what downstream DE analysis results may look like (e.g. poor separation between biological groups in the MDS plot tends to result in no or low numbers of differentially expressed genes).

#### Sizing, shaping and colouring points

Typically, one would simply examine the first 2 dimensions in a static MDS plot - the first 2 dimensions explain most of the variation in the data; and move onto the next steps in data exploration if satisfied with the clustering of biological and/or experimental groups there. Whilst this task is fairly straight-forward, one can easily end up with several versions (4 or more) of the same dimension-1-and-2 MDS plot by the time they apply different sizing, shapes and colours to the plot to reflect experimental factors in the study. Applying these custom visual transformations to static R plots can be time-intensive and require coding experience. We therefore designed dashboard-style inputs which apply a range of visual modifications to the MDS plot. Users can encode information about each sample by adjusting the colour, shape and size of each point using drop-down menus (see Figure 2). Points can be coloured by a given feature using a continuous or discrete colour scheme of choice which can be changed on the fly using a drop-down menu. These modifications to the MDS plot help users determine the experimental factors by which samples cluster.

They also allow for the encoding of redundancy into a visualisation which can emphasise the visual groupings of a data display (24). For instance in Figure 2, samples are coloured and shaped by the same experimental factor, whilst library sizes are reflected in its sizing.

#### Plotting Dimensions

Aside from dimension-1-and-2 plots, the examination of higher dimensions allows one to check for unwanted batch effects in the dataset (13). In many cases, this next step is skipped over if dimension-1-and-2 plots do not raise any questions, especially when production of the first set of plots was already quite time-intensive. Occasionally, one would create higher dimension R plots within the R console as a visual inspection, but not save or document the plot unless an interesting feature is discovered (e.g. samples cluster by sample collection date when sample collection date was not expected to influence gene expression). *Glimma* 1.0 only allowed the user to plot consecutive pairs of dimensions together (i.e. 1 and 2 in conjunction, 2 and 3, etc). Version 2.0 remedies this limitation by providing drop-down menus which allow the user to plot any combination of dimensions on the x and y axes. This allows the creation of plots that show separation of samples based on two experimental factors when there is a confounding factor driving an intermediate MDS dimension. Moreover, the interactive plots allow one to check variations of the plot (higher dimensions or combination of dimensions) without having to retrospectively code and save extra plots to an analysis. The new “Save Plot” button (see Figure 2) is also useful for adding any variation of the MDS plots to reports, presentations and publications when the interactive form is not needed or when working outside of R Markdown documents.

### Summary-Expression Plot Improvements

Summary-expression plots in *Glimma*, which include MA, volcano and XY plots, share a common front-end user interface (see Figure 3), with a main plot of gene-wise summary statistics on the top-left and an expression plot showing the abundance of the selected gene in each sample on the top-right. Beneath these plots lies a table containing annotation data and DE analysis statistics which is interactively linked to both the summary and expression plots.

The MA plot, which is called via the `glimmaMA` function, is used to visualise logFC (M) on the y-axis versus average abundance (A) on the x-axis. The volcano plot, called via the `glimmaVolcano` function, displays a measure of statistical significance on the y-axis, specifically this is the -log_10_-*p*-value of genes where higher values reflect greater significance, versus logFC on the x-axis. The generic XY plot, which is called via `glimmaXY`, displays two userparameterised vectors on the x-and y-axes of the summary plot. The vectors are required to be same length as the number of genes in the dataset, such that the points in the summary plot must match the number of columns in the expression data.

Numerous improvements have been made to the R function interfaces and common front-end for this trio of summary-expression plots. These are summarised below.

#### Volcano Plot

In previous versions of *Glimma*, volcano plots could only be generated by manually extracting and log-transforming the relevant x and y axis vectors, and then passing these to the `glimmaXY()` function. Version 2.0 provides a new `glimmaVolcano` function that automatically extracts and transforms statistical significance alongside logFC from *edgeR, limma* and *DESeq2* objects to generate an interactive volcano plot widget as per Figure 3.

#### Multiple Gene Selections

In DE analysis we are often interested in the transcription profile of a set of multiple genes. In the initial release of *Glimma*, researchers had to query genes one-by-one in order to locate them on the plot. Version 2.0 remedies this by allowing for multiple genes to be simultaneously highlighted. Users can search and sort genes in the table of summary statistics and annotation data. Selecting any number of rows on the table will highlight the corresponding points on the graph. For instance, the three most significant genes on the X chromosome have been highlighted in Figure 3. Interaction is bi-directional: users can also click on points in the summary plot to toggle their selection, which will filter the table beneath to display data for just the selected genes. The expression plot on the right shows the expression values measured in log_2_-counts-per-million (log-CPM) across all samples for the most recently selected gene, stratified by experimental group.

#### Automatic Highlighting of Significant Genes

Version 2.0 auto-highlights significant genes on behalf of the user by calling `limma::decideTests()` on `MArrayLM` objects and `edgeR::decideTestsDGE()` on `DGEExact` and `DGELRT` objects. These sensible defaults can be overridden using the `status` argument.

#### User-Specified Colours for Expression Plot

Summary-expression plots take an optional `sample.cols` vector argument which allows users to vary the colours of samples in the expression plot on. This creates a visualisation whereby samples are separated by treatment group but coloured by another factor such as gender or batch.

#### X-axis Label Rotation

Feedback from version 1.0 users suggested that when a large number of groups were plotted on the expression plot, distinguishing between the labels of each group became impossible. Version 2 allows a larger number of clearly readable x-axis labels by rotating each label by 45 degrees anti-clockwise.

#### Fixing Expression Y-axis

By default, selecting a gene causes the vertical axis of the expression plot to re-scale itself according to the maximum expression value for that gene. Feedback from biologists indicated that this made it difficult to notice when there were large changes in expression relative to another gene they had clicked on. Version 2.0 remedies this by adding the `max_y_axis` input form allowing the user to fix the y-axis maximum to the numeric value specified (see Figure 3), which overrides the auto-scaling behaviour.

#### User Interface Improvements

Plots now display gene symbol text above selected points, allowing genes to be visually identified without additional manipulation in Illustrator. Legends were added to the summary plot indicating the colour associated with the DE status of the gene (1 for upregulated, 0 for non-DE, -1 for downregulated). A list of gene identifiers beneath the plots indicates the current selection of genes highlighted across the entire table or plot, helping the user to keep track of what they have highlighted.

#### Data Export

The “Save Data” button allows researchers to export the table data concatenated with expression data as a CSV file. Researchers can optionally save only the selected subset of genes, or all genes in the table. This is useful for further interrogation of the data using external programs and/or manipulation into other plotting styles outside of *Glimma*.

#### Streamlined Function Arguments

Plots in version 1.0 had lengthy function prototypes that required the user to manually extract gene counts, test results and experimental groups from data structures which already contained this information. Version 2.0 significantly reduces the verbosity of function calls by automating data extraction for the user. For instance, function calls for the `MArrayLM` data structure from *limma* now require two arguments down from four:

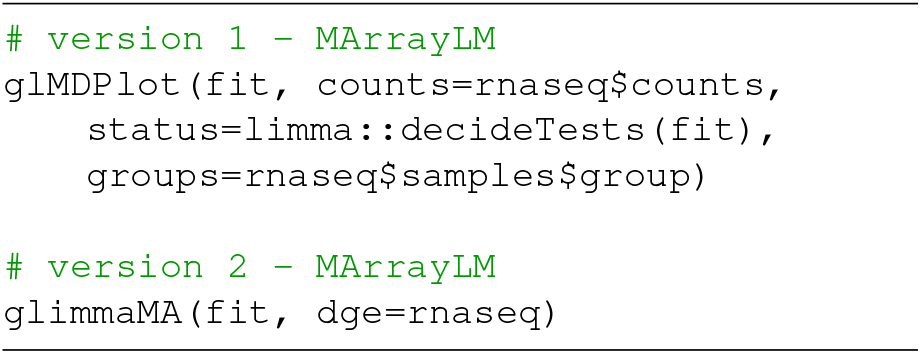

In a similar fashion, analysis plot function prototypes for *edgeR* and *DESeq2* data structures now require 2-3 fewer arguments to generate the equivalent plots.

### Plots for scRNA-seq data

The *Glimma* package was originally designed and developed with bulk RNA-seq data in mind. However, we have found that users are also applying the plots to scRNA-seq data. In light of this, we have provided preliminary support for single cells as a new vignette to demonstrate the use of *Glimma* version 2.0 for single cell data. Briefly, MDS plots can be created using the full set of single cells, while summary-expression plots can be created for pseudo-bulk samples or a sub-sample of the single cells. Sub-sampling of the single cells is currently necessary to meet performance constraints, but a random sub-sample is still informative to the distribution of the full set of samples. Rotation of x-axis labels has also improved the applicability of *Glimma* to scRNA-seq analyses which often contain a high number of sample groups or cell types.

## DISCUSSION

In this paper we presented R markdown-ready interactive plots for gene expression analysis, designed for convenient use for biologists and end users. A highly requested feature pack has been implemented which improves the reproducibility and interactivity of the package, with major inroads also made to supporting single cell RNA-seq analysis. Reimplementing *Glimma* using high-level libraries has greatly increased the extensibility of the package, future-proofing it for new capabilities to be added rapidly in response to user demand. For instance, version 2 achieved a 63.8% reduction in lines of R code (1,175 versus 3,246 in version 1.0) despite incorporating many more features due to the use of external libraries and modularisation. This translates into a vastly more manageable codebase.

Another noteworthy concern is the file size of knitted HTML from R markdown containing *Glimma* 2.0 interactive plots. For the plots demonstrated in this paper drawn from a dataset with nine samples and 27,000 genes, we recorded file sizes of 1.1 and 4.5 megabytes for the MDS and volcano plots respectively. As we expect file sizes to increase linearly with the cardinality of the datasets used in analysis, compressing exported plots may be prudent in order to share them around more efficiently.

A Bioconductor tool that shares similar goals to *Glimma* is the *iSEE* package (9) which allows users to dynamically configure data displays such that one panel transmits certain features of the data to another receiving panel. For instance, panels can be arranged such that selecting rows in a transmitting table filters points in a receiving graph, achieving a similar effect to the *Glimma* analysis plots. In *Glimma*, transmission and reception of data between displays is pre-configured and immutable, sacrificing flexibility for ease of use and minimal set-up.

### Future Work

*Glimma*’s interactive analysis plots can be prohibitively slow to use on scRNA-seq analyses. This is due to the size of scRNA-seq counts matrices which can contain many thousands of samples. Taking advantage of the fact that the vast majority of counts are zero in single cell expression data, we are exploring sparse matrix objects for JavaScript to improve performance. The main challenge in this space is selecting and testing an open-source sparse matrix implementation which maximises the efficiency for our use case (storing scRNA-seq data) while adding minimal bulk to the package.

In addition, the direct support of `SingleCellExperiments` objects from Bioconductor is challenging due to the lack of consensus among different packages on how final summary statistics are to be stored. Direct support of `SingleCellExperiment` objects can be added when a consensus or dominant method emerges.

In future versions of *Glimma*, we aim to extend the interactive front-end for the MDS plot to support UMAP (25) and tSNE (26) dimensionality reduction visualisations. Further improvements to reproducibility of the MDS plot could also be achieved by allowing users to pre-set the initial colour, shape or size of points using function arguments.

## ACKNOWLEDGEMENTS

We thank Stuart Lee, Peter Hickey, Luyi Tian, Monther Alhamdoosh, Emma Gail and Iromi Wanigasuriya for their helpful feedback on this work.

## Funding

CWL was supported by the Chan Zuckerberg Initiative Essential Open Source Software for Science Program (Grant no. 2019-207283) and by CSL Limited.

## Conflict of interest statement

None declared.

## Notes

### Competing Interest Statement

The authors have declared no competing interest.

## References

1. Zhong Wang, Mark Gerstein, and Michael Snyder. RNA-Seq: a revolutionary tool for transcriptomics. Nature Reviews Genetics, 10(1):57–63, January 2009. ISSN 1471-0064. doi: 10.1038/nrg2484.x

2. R Core Team. R: A Language and Environment for Statistical Computing. R Foundation for Statistical Computing, Vienna, Austria, 2020. URL: https://www.R-project.org/.

3. S. Su, C. W. Law, C. Ah-Cann, M.-L. Asselin-Labat, M. E. Blewitt, and M. E. Ritchie. Glimma: interactive graphics for gene expression analysis. Bioinformatics, 33:2050–2052, 2017.

4. M. E. Ritchie, B. Phipson, D. Wu, Y. Hu, C. W. Law, W. Shi, and G. K. Smyth. limma powers differential expression analyses for RNA-sequencing and microarray studies. Nucleic Acids Res, 43(7):e47, 2015.

5. M. D. Robinson, D. J. McCarthy, and G. K. Smyth. edgeR: a Bioconductor package for differential expression analysis of digital gene expression data. Bioinformatics, 26:139–40, 2010.

6. Michael I. Love, Wolfgang Huber, and Simon Anders. Moderated estimation of fold change and dispersion for RNA-seq data with DESeq2. Genome Biology, 15(12):550, December 2014. ISSN 1474-760X. doi: 10.1186/s13059-014-0550-8.

7. Alper Kucukural, Onur Yukselen, Deniz M. Ozata, Melissa J. Moore, and Manuel Garber. DEBrowser: interactive differential expression analysis and visualization tool for count data. BMC Genomics, 20(1):6, January 2019. ISSN 1471-2164. doi: 10.1186/s12864-018-5362-x.

8. Federico Marini, Jan Linke, and Harald Binder. ideal: an R/Bioconductor package for interactive differential expression analysis. BMC Bioinformatics, 21(1):1–16, December 2020. ISSN 1471-2105. doi: 10.1186/s12859-020-03819-5.

9. Kevin Rue-Albrecht, Federico Marini, Charlotte Soneson, and Aaron T.L. Lun. iSEE: Interactive SummarizedExperiment Explorer. F1000Research, 7:741, June 2018. ISSN 2046-1402. doi: 10.12688/f1000research.14966.1.

10. Jonathan W Nelson, Jiri Sklenar, Anthony P Barnes, and Jessica Minnier. The START App: a web-based RNAseq analysis and visualization resource. Bioinformatics, 33(3):447–449, February 2017. ISSN 1367-4803. doi: 10.1093/bioinformatics/btw624.

11. Adam Price, Adrian Caciula, Cheng Guo, Bohyun Lee, Juliet Morrison, Angela Rasmussen, W. Ian Lipkin, and Komal Jain. DEvis: an r package for aggregation and visualization of differential expression data. 20(1):110. ISSN 1471-2105. doi: 10.1186/s12859-019-2702-z.

12. Winston Chang, Joe Cheng, J. J. Allaire, Yihui Xie, and Jonathan McPherson. shiny: Web Application Framework for R, June 2020. URL: https://CRAN.R-project.org/package=shiny.

13. C. W. Law, M. Alhamdoosh, S. Su, X. Dong, L. Tian, G. K. Smyth, and M. E. Ritchie. RNA-seq analysis is easy as 1-2-3 with limma, glimma and edgeR. F1000Research, 5(1408), 2016.

14. Michael Bostock, Vadim Ogievetsky, and Jeffrey Heer. D3: Data-Driven Documents. IEEE Transactions on Visualization and Computer Graphics, 17(12):2301–2309, December 2011. ISSN 1077-2626. doi: 10.1109/TVCG.2011.185.

15. Ramnath Vaidyanathan, Yihui Xie, J. J. Allaire, Joe Cheng, Carson Sievert, Kenton Russell, and Ellis Hughes. htmlwidgets: HTML Widgets for R, December 2020. URL: https://CRAN.R-project.org/package=htmlwidgets.

16. Arvind Satyanarayan, Ryan Russell, Jane Hoffswell, and J. Heer. Reactive Vega: A Streaming Dataflow Architecture for Declarative Interactive Visualization. IEEE Transactions on Visualization and Computer Graphics, 2016. doi: 10.1109/TVCG.2015.2467091.

17. Yihui Xie, J.J. Allaire, and Garrett Grolemund. R Markdown: The Definitive Guide. Chapman and Hall/CRC, Boca Raton, Florida, 2018. ISBN 9781138359338.

18. Wolfgang Huber, Vincent J. Carey, Robert Gentleman, Simon Anders, Marc Carlson, Benilton S. Carvalho, Hector Corrada Bravo, Sean Davis, Laurent Gatto, Thomas Girke, Raphael Gottardo, Florian Hahne, Kasper D. Hansen, Rafael A. Irizarry, Michael Lawrence, Michael I. Love, James MacDonald, Valerie Obenchain, Andrzej K. Oleś, Hervé Pagès, Alejandro Reyes, Paul Shannon, Gordon K. Smyth, Dan Tenenbaum, Levi Waldron, and Martin Morgan. Orchestrating high-throughput genomic analysis with Bioconductor. Nature Methods, 12(2):115–121, February 2015. ISSN 1548-7105. doi: 10.1038/nmeth.3252.

19. Carson Sievert. Interactive Web-Based Data Visualization with R, plotly, and shiny. Chapman and Hall/CRC, 2020. ISBN 9781138331457. URL: https://plotly-r.com.

20. Dan Vanderkam, Petr Shevtsov, J. J. Allaire, RStudio, Jonathan Owen, Daniel Gromer, Benoit Thieurmel, Kent Laukhuf, jQuery Foundation, and jQuery contributors. dygraphs: Interface to ‘dygraphs’ interactive time series charting library. URL: https://CRAN.R-project.org/package=dygraphs.

21. Eli Grey. FileSaver.js, December 2020. URL: https://github.com/eligrey/FileSaver.js.

22. Julie M. Sheridan, Matthew E. Ritchie, Sarah A. Best, Kun Jiang, Tamara J. Beck, François Vaillant, Kevin Liu, Ross A. Dickins, Gordon K. Smyth, Geoffrey J. Lindeman, and Jane E. Visvader. A pooled shRNA screen for regulators of primary mammary stem and progenitor cells identifies roles for Asap1 and Prox1. BMC Cancer, 15(1):221, April 2015. ISSN 1471-2407. doi: 10.1186/s12885-015-1187-z.

23. Y. Liao, G. K. Smyth, and W. Shi. The Subread aligner: fast, accurate and scalable read mapping by seed-and-vote. Nucleic Acids Res, 41:e108, 2013.

24. Christine Nothelfer, Michael Gleicher, and Steven Franconeri. Redundant encoding strengthens segmentation and grouping in visual displays of data. Journal of Experimental Psychology. Human Perception and Performance, 43(9):1667–1676, September 2017. ISSN 1939-1277. doi: 10.1037/xhp0000314.

25. Leland McInnes, John Healy, and James Melville. UMAP: Uniform Manifold Approximation and Projection for Dimension Reduction. 1802.03426 [cs, stat], September 2020. arXiv: 1802.03426.

26. Laurens van der Maaten and Geoffrey Hinton. Visualizing data using t-SNE. Journal of Machine Learning Research, 9:2579–2605, 2008.

